# Optimizing an eDNA protocol for monitoring endangered Chinook Salmon in the San Francisco Estuary: balancing sensitivity, cost and time

**DOI:** 10.1101/871368

**Authors:** Thiago M. Sanches, Andrea M. Schreier

**Affiliations:** Animal Sciences, University of California-Davis, Davis, California, US

## Abstract

Environmental DNA (eDNA) analysis has gained traction as a precise and cost effective method for species and waterways management. To date, publications on eDNA protocol optimization have focused primarily on DNA yield. Therefore, it has not been possible to evaluate the cost and speed of specific components of the eDNA protocol, such as water filtration and DNA extraction method when designing or choosing an eDNA pipeline. At the same time, these two parameters are essential for the experimental design of a project. Here we evaluate and rank different eDNA protocols in the context of Chinook salmon (*Oncorhynchus tshawytscha*) eDNA detection in an aquatic environment, the San Francisco Estuary. We present a comprehensive evaluation of multiple eDNA protocol parameters, balancing time, cost and DNA yield. For estuarine waters, which are challenging for eDNA studies due to high turbidity, variable salinity, and the presence of PCR inhibitors, we find that a protocol combining glass filters and magnetic beads, along with an extra step for PCR inhibitor removal, is the method that best balances time, cost, and yield. In addition, we provide a generalized decision tree for determining the optimal eDNA protocol for other studies on aquatic systems. Our findings should be applicable to most aquatic environments and provide a clear guide for determining which eDNA pipeline should be used for a given environmental condition.

**Author Summary:** The use of environmental DNA (eDNA) analysis for monitoring wildlife has steadily grown in recent years. Though, due to differences in the ecology of the environment studied and the novelty of the technique, eDNA currently shows a lack of standards compared to other fields. Here we take a deep look into each step of an eDNA assay, looking at common protocols and comparing their efficiencies in terms of time to process the samples, cost and how much DNA is recovered. We then analyze the data to provide a concise interpretation of best practices given different project constraints. For the conditions of the San Francisco Estuary we suggest the use of glass fiber filtration, the use of paramagnetic beads for DNA extraction and the use of a secondary inhibitor removal. We expect our findings to provide better support for managers to decide their standards ahead of project submission not only for estuarine conditions but for other waterine conditions alike.

## Introduction

Environmental management policies rely heavily on measurements of the spatial distribution of habitat occupancy of species. In the past decade, environmental DNA (eDNA) has gained traction as one of the most sensitive and cost effective monitoring methods [1], allowing researchers to better estimate species occupancy rates in a given habitat. Due to high variability in the studied environments, currently, there are no clear guidelines to assist investigators in choosing an optimal protocol for their particular eDNA monitoring studies.

In this study, we separate and optimize four important steps for eDNA biomonitoring of delta estuarine waters, which are characterized by elevated concentrations of solid suspended particles and fluctuating levels of salinity [2]. We comment on the specifics of each step for eDNA biomonitoring and develop a concise guideline to help determine which approach is most suitable for a set of scenarios. To lessen the burden of comparing DNA isolation methods on future investigators, we provide a framework that should help make more informed decisions, taking into account the specifics of their study requirements.

### The four main steps of an eDNA pipeline

The protocol to isolate environmental DNA from water samples can be described in four steps: filtration, DNA extraction, inhibitor removal and DNA amplification in order to estimate the initial concentration of eDNA [3]. In the filtration step, the water samples, with preferred volumes ranging from 50 mL to 1 liter in previous studies, are pressure-pumped through a membrane filter which captures the free DNA as well as tissue and cells suspended in the water. The next step is to extract DNA from the filter using conventional extraction methods, which were developed to isolate large nuclear DNA fragments from tissue. However, in the case of eDNA, it is preferable to target small fragments of mitochondrial genes as they have a higher copy number per cell compared to nuclear DNA. Then, as the filter may also capture high concentrations of PCR inhibitors, it is often necessary to use a secondary inhibitor removal step to further isolate the DNA from contaminants [4]. Last, the isolated DNA is amplified using quantitative PCR (qPCR) with primers specific to the target species, and the initial amount of the target eDNA is determined based on the Cq method. Although it is possible to use conventional PCR, this method significantly underperforms compared to qPCR in terms of sensitivity, and even when DNA is successfully amplified it provides less informative data [5].

### PCR inhibitors

PCR inhibitors are a group of substances that can inhibit PCR amplification. Their inhibiting mechanism varies between affecting the template DNA, the polymerase or other reagents necessary for the reaction. PCR inhibitors can be catalytic (e.g. proteases degrading proteins and phenol degrading DNA) or work through competitive binding (e.g. melanin forming a complex with the polymerase and humic acid interacting with the DNA template) [6]. Humic matter and proteases are typical PCR inhibitors present in high concentrations in turbid waters and other environmental samples [6,7].

### Estuarine waters and fish detection

In this study we optimize eDNA biomonitoring for estuarine waters, as this habitat is essential for the early developmental stages of several anadromous species, including our target species, Chinook salmon (*Oncorhynchus tshawytscha*). The estuarine environment provides a challenge for eDNA biomonitoring as the elevated density of solid particles, measured by turbidity levels, can bind to the DNA and clog the pores of the filters, limiting the volume of water that might be filtered. Also, estuaries have been shown to have elevated levels of PCR inhibitors [8,9]. Therefore, we assume that if our DNA amplification-based experiments work in these complex conditions, the same approach could also be applied to less turbid freshwater and marine conditions.

### Chinook Salmon as a target

We targeted Chinook Salmon in our experiments for a variety of reasons. First, as a widespread species in the North American Pacific Northwest, it has invaluable importance for the stability of the marine ecosystem of the region [10] and at the same time, provides a critical source of income for historic fishing communities [11,12]. In California, the Central Valley Spring-run, Fall-run and Winter-run Chinook salmon are listed as vulnerable while the Sacramento River winter-run and Central Valley late Fall-run are listed as endangered [13]. Little is yet known about the spatio-temporal distribution and estuarine habitat usage of pre-smolt juvenile Chinooks during their annual out migration to the ocean. In this life-stage, juvenile Chinook salmons runs blend together and use this period to grow before leaving the estuarine environment. Larval survival in this stage has major implications to the population size of the species [14]. Conventional survey methods have encountered numerous difficulties to sample the rare juvenile Chinook in the marshland conditions of the SFE. Developing a high precision, high throughput eDNA protocol optimized for estuarine waters will allow managers to have a better understanding of the habitats used by Chinook in their early life-stages.

### Accounting for cost and time in experimental design

To decide on the most practical estuarine eDNA protocol, we first need to determine what it means for a protocol to be efficient. In our case we listed our priorities in the following order: 1) The eDNA yield must be adequately sensitive in realistic scenarios; 2) The protocol must be fast and scalable, and 3) The protocol must be cost effective, considering that reagent cost is the main driver of cost per sample. This order of priorities is influenced by several factors that include species abundance and costs. If the target species is ubiquitous and present at high densities, the DNA yield constraint can be loosened, allowing the use of faster and more cost-conscious protocols. If labor cost is inexpensive, choosing a more time intensive yet cheaper protocol will maximize the number of sampling points. On the other hand, in situations where labor accounts for much of the costs, choosing less time intensive protocols will allow more sampling points for the project.

## Results and Discussion

### Interference between filter type and extraction method is negligible

We first examined different methods for the initial two steps of an eDNA pipeline, filtration and DNA extraction, and tested for interference between these steps. In general, when optimizing a protocol consisting of several steps, it is important to identify if previous steps interfere with the effectiveness of subsequent steps. In our case the main possible interference is between the filter used and the extraction method. The yield percentage of a certain extraction method could change depending on which filter was used. Possible reasons for interference between filter and extraction method include different particles binding differentially to filters and extraction methods not isolating DNA from all types of particles at the same yield percentage. The models that we tested are the following:

Model with interference:

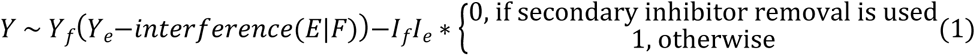

Model without interference:

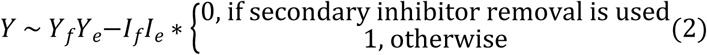

For both widely-applicable information criterion (WAIC) and Leave-one-out cross validation (LOO) the model without interference was selected with a weight of 1 in both cases [17,20]. One form of visualizing the absence of interactions is that the ranking order of filter yield (1st *Cellulose nitrate - 2nd Glass fiber - 3rd Filter paper N1)* does not change independently of which extraction method is chosen (Fig 1A). Similarly, the yield ranking for extraction methods is not affected by filter choice. (Fig 1B). Only the NaOH method breaks the independence rule for the nitrocellulose filter. In this case, target DNA could not be amplified from NaOH extractions without secondary inhibitor removal, resulting in an upwards skewed average of the DNA yield as the samples without secondary inhibitor removal were not taken into account.

**Figure 1:**
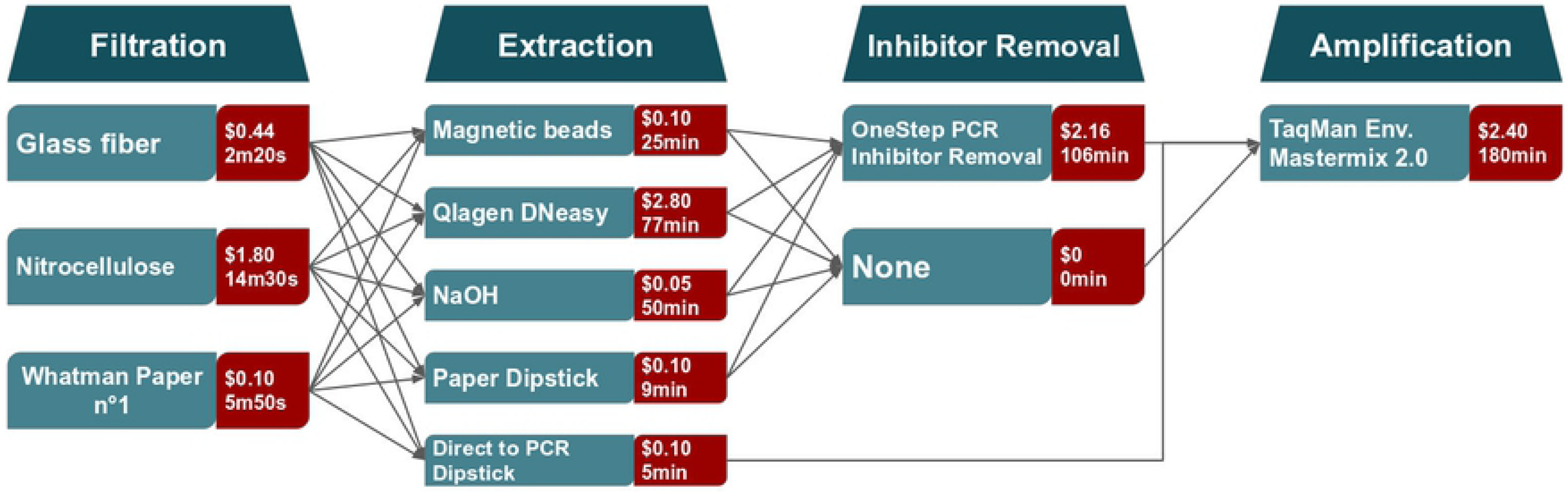
Relationship between total DNA yield and filtration and extraction protocols. (A) No crossing between lines indicates that on average, DNA yield ranking for the filters is independent of the extraction protocol. Error bars represent the 95% confidence intervals. The combination of Whatman filter and NaOH extraction wasn’t able to amplify the target Chinook DNA. (B) No crossing between lines indicates that on average, DNA yield ranking for the extraction method is independent of the filter type. The NaOH extraction protocol is the only case where the ranking order is not maintained and can be explained by the added effects of carry-on inhibitors.

The lack of interference between steps shows that for future optimization tests, it is not necessary to test all the possible combinations of filters and extraction methods at the same time. Instead one might test each section of the protocol independently and converge on the optimal method. Therefore, it is possible to test more methods for each step and increase the number of replicates for each test in future optimization experiments.

### The nitrocellulose filter can retain the most DNA per volume while the glass filter is the most resilient to high levels of turbidity

Next, we compared DNA yields from three different filters. The nitrocellulose filter outperformed the glass fiber filter in terms of DNA yield by 1.6 times and the Whatman n°1 filter by 3.75 times on average (Fig 2). In other words, 1.6L and 3.75L of water would need to be filtered through a glass fiber filter or Whatman n°1 filter respectively to isolate the same amount of DNA as filtering 1L of water through a nitrocellulose filter. However, the glass filter outperforms the nitrocellulose and Whatman filters in terms of filtration time, with the glass filter not only being drastically faster but also more consistent and resilient to variations in turbidity (Fig 3). Therefore, we can conclude that for the context of estuarine waters, the glass filter is optimal in terms of DNA yield, speed, and cost.

**Figure 2:**
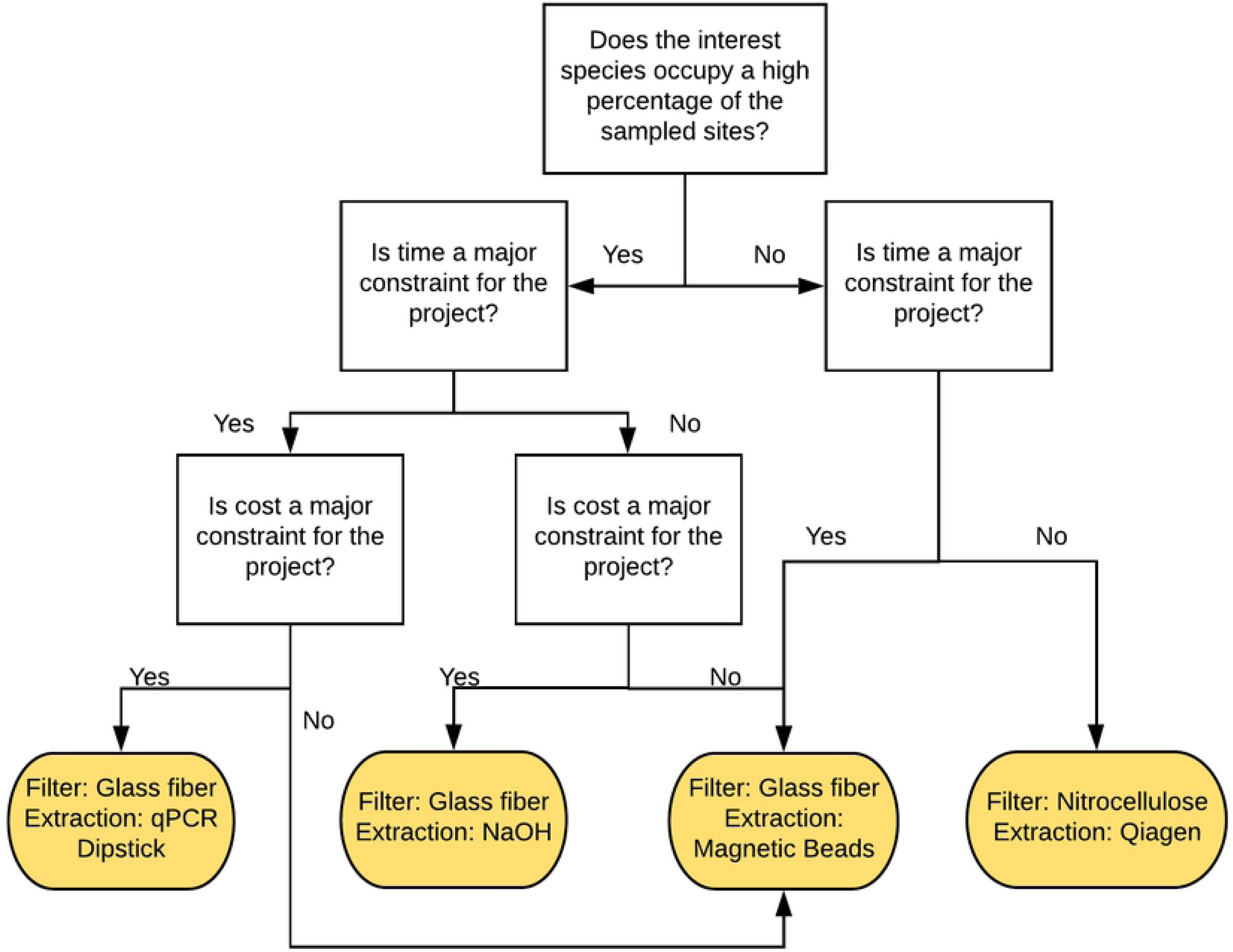
Distribution of DNA capture ratio for each filter type from the Automatic Differentiation Variational Inference model. The broadness of the curve shows the variability of the ratio of the input DNA that binds to the filter. The peak of each distribution is the mean yield ratio of DNA recovery for that filter type. The nitrocellulose filter yielded the highest recovery ratio with little efficiency overlap compared to glass and Whatman filters.

**Figure 3:**
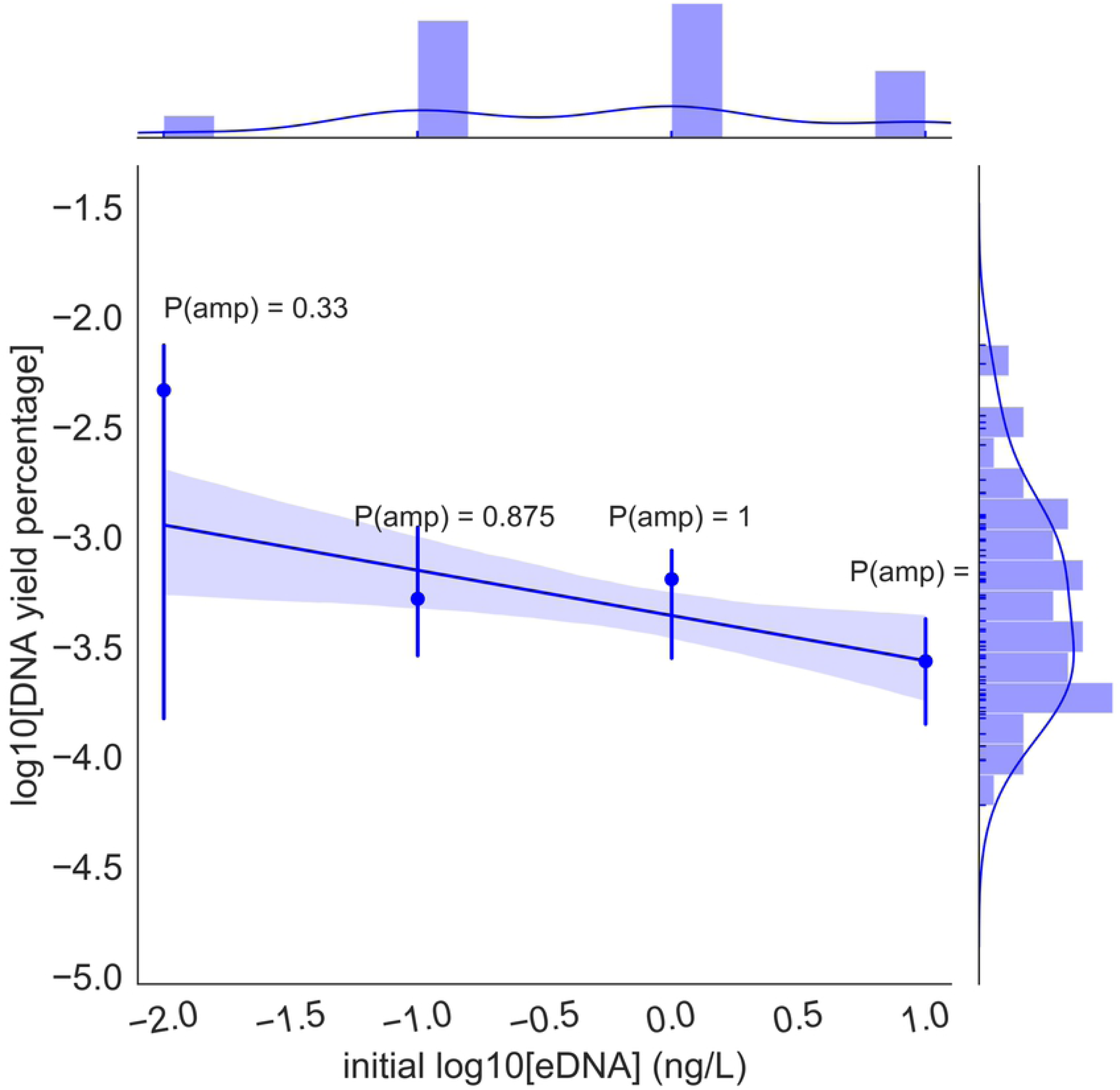
Percentiles and medium filtration time in order to filter 1L of estuarine water for each filtration method. The glass filter outperforms nitrocellulose and Whatman by a significant margin in terms of average filtering time and consistency in the filtering time. Dots are filtration events while the black line represents the median value filtering time. Boxes indicate 10% quantiles.

### QIagen DNA extraction is the most sensitive and reliable, paper extraction is the fastest and most cost effective, and magnetic beads is the most balanced method

All extraction methods could yield enough eDNA to be detectable by qPCR amplification. The Qiagen DNEasy kit had the highest DNA yield, outperforming NaOH by 1.7 times, magnetic beads by 2.26 times, direct to qPCR dipsticks by 9.71 times and regular dipsticks by 358 times (Fig 4). At the same time, the Qiagen kit is by a considerable margin the most time-consuming method, requiring 77 minutes to process 18 samples. In contrast, the direct to qPCR dipstick approach was the fastest and most cost-efficient method by a wide margin. Currently, the major bottleneck of our experiments is the time required to process the samples. Yet, subsequent tests have shown that the use direct dipstick extraction drastically lower the probability of amplification in cases where the species of interest is rare. Therefore, we consider the magnetic beads to be the optimal method for estuarine waters.

**Figure 4:**
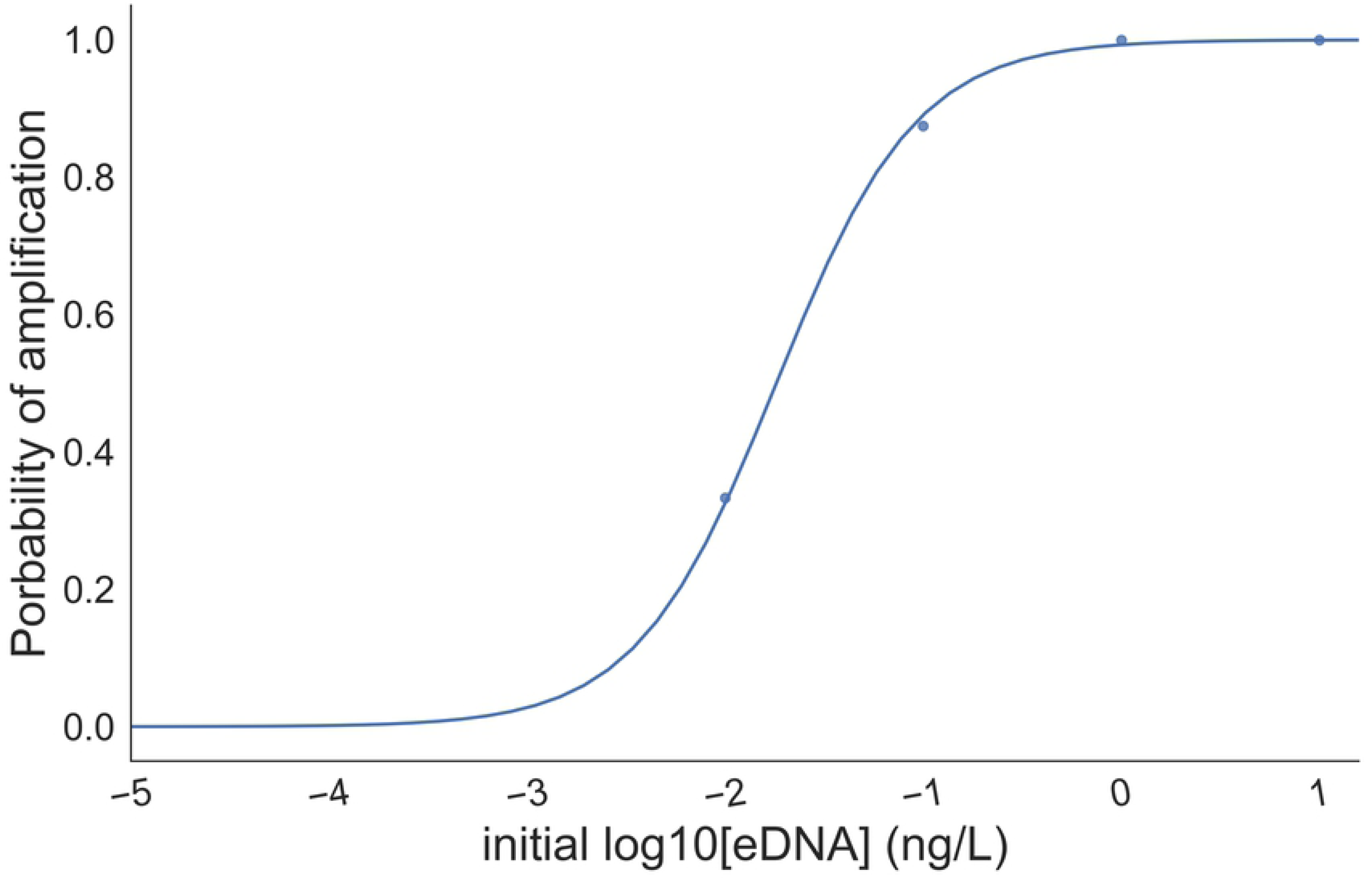
Modelled distribution of percentage yield for each extraction protocol. The width of each of the curves shows the variability in modelled yield. The peak of the distribution is the mean yield per extraction type. QIagen DNeasy yields the best yield with little overlap with other methods. Meanwhile magnetic beads and NaOH have shown similar distributions with significant overlap, while both dipstick methods underperform the other methods. It is important to note that this plot does not take into account PCR inhibitor carryover, which might vary significantly between methods.

We estimated our costs for the most-used DNA extraction kits. Alternative kits might be used in order to reduce costs. As an example, Ampure XP is 100 times more expensive than making a magnetic beads solution in-house [21], though this cost reduction is at the expense of lower reproducibility and therefore not optimal for certain projects. Buying in bulk is also other alternative to reduce costs, though that might be limited to initial funding of the project.

### Yield is mostly dependent on extraction method

The extraction method was shown to be the most influential factor for the eDNA yield from the random forest aggressor analysis (Fig 5). Therefore, further optimization experiments should focus on this step, experimenting with different protocols to extract the eDNA in order to maximize protocol eDNA yield. Meanwhile, in the context of our experiments, the removal of inhibitors was shown to have little impact to the total DNA yield estimated by qPCR, although published data [4] have shown that inhibitor removal highly influences the amplification probability of the qPCR reaction.

**Figure 5:**
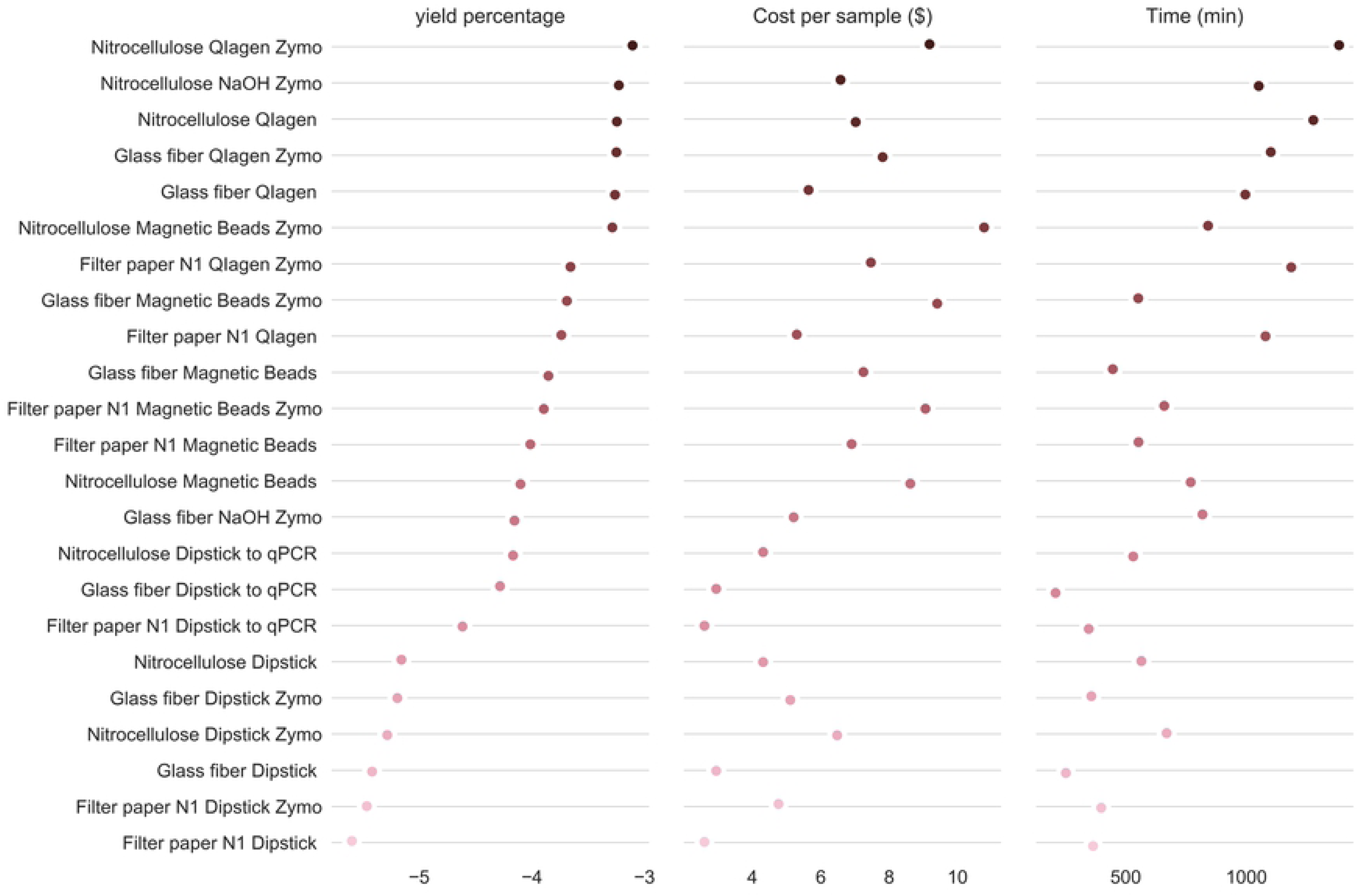
Influence of each eDNA protocol segment to total eDNA yield ratio estimate. The extraction method was the factor that had the highest influence on the total eDNA yield of the total protocol, while inhibitors didn’t have a significant impact compared to the other sections of the total protocol.

Our experiments have shown comparatively little influence of the use of secondary inhibitor removal to the total eDNA yield. Though this result contradicts at some degree our pilot study for estuarine juvenile chinook as well as other published data. Causes for this discrepancy might be resultant of several nonexclusive factors. First inhibitors in general work by binding to DNA strands and not as catalysts, therefore if the ratio between eDNA:inhibitors is significantly elevated, as we would expect in tank experiments, we would expect minimal effects of the inhibitors. Another possibility is that at the sampling location, on the sampling date and time, there were fewer inhibitors than usually observed in estuaries. Another possible explanation is that the yield variance between the extraction methods and filters surpass the yield variance due to the inhibitor removal step, which doesn’t mean this step won’t significantly influence the DNA yield of the experiment. Last, PCR inhibitors might not affect the eDNA retrieval but only the probability of amplification. This last observation might also explain why the probability of amplification and DNA yield aren’t always fully correlated. Therefore, considering this experiment results and previous findings we consider that secondary inhibitor removal is advised if possible as it improves the DNA yield and amplification probability in the context of estuarine samples.

### Effects of secondary inhibitor removal varies between filters and extraction method

We observed that the secondary inhibitor removal step always outperformed skipping this step. Regressions from Figs 6A and 6C were always positive and the distributions from Figs 6B and 6D were always greater than zero. The nitrocellulose filter and the NaOH extraction were the methods that carried the most PCR inhibitors, while the other methods for each step showed a high overlap of their carryover inhibitor distributions. Secondary inhibitor removal was essential to observe any amplification using the NaOH extraction method, which also suggested that this method is inefficient at removing PCR inhibitors.

**Figure 6:**
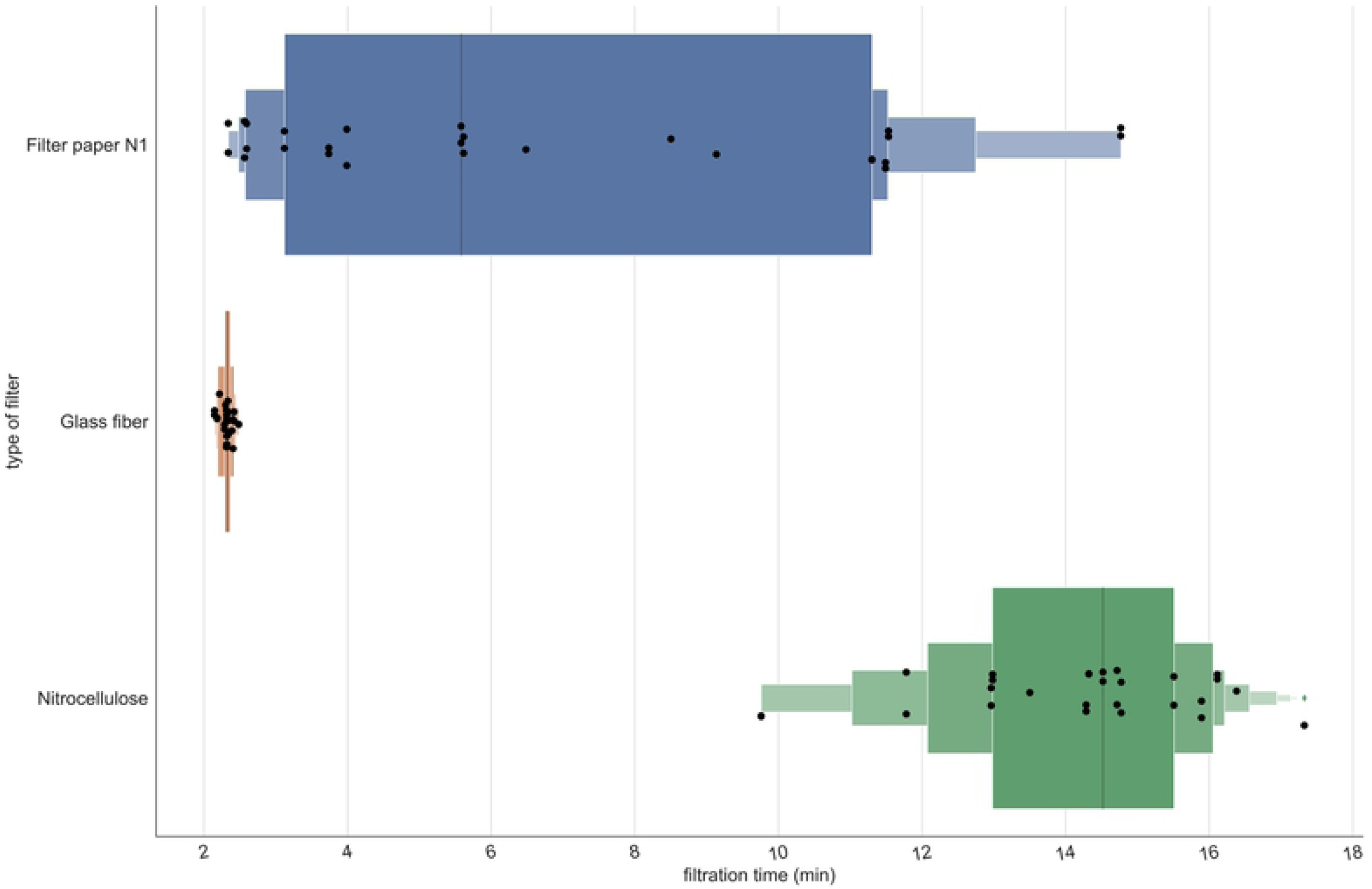
Effects of adding a secondary inhibitor removal step to the eDNA estimation protocol. (A-B) DNA yield variation of using a OneStep PCR Inhibitor Removal^©^. The nitrocellulose filter produced the highest inhibitor carryover levels at the same time it captured the highest percentage of free eDNA. This suggested that the nitrocellulose filter captured particulates with indiscriminately with high efficiency (C-D) Estimated distributions for inhibitor carryover for filter and extraction method. Aside from NaOH extraction, other methods had similar distributions of carryover PCR inhibitors with high overlap. Therefore, NaOH extraction, even if it has an elevated eDNA yield, doesn’t properly address the high levels of PCR inhibitors commonly encountered in environmental samples.

In most cases, using a glass fiber filter and magnetic beads would be the most practical method to generate the maximum amount of information obtained about fish distribution given the constraints of our study. Our experiment suggests that DNA extraction from the filters is the most time-consuming step and most variable in terms of efficiency; therefore, this is the step which should be decided with utmost care in order to maintain the high-throughput and useful detection limit of the desired methodology. For this reason, magnetic beads DNA extraction in a promising alternative to silica column extraction, as this method strikes the balance between yield, amplification probability, carryover PCR inhibitors and time to process samples. Meanwhile the cost of using magnetic beads can be mitigated by developing the necessary reagents in-house. Though in specific cases different pipelines might yield better results. For those scenarios, we constructed a simple decision tree for choosing the best methodology various possible study for each scenario (Fig 7). We also ranked the pipelines, sorting them by DNA yield, which should be the main parameter for the pipeline selection. Then, once established which pipelines have a DNA yield that fits the project, balancing time and cost of the pipelines (Fig 8).

**Figure 7:**
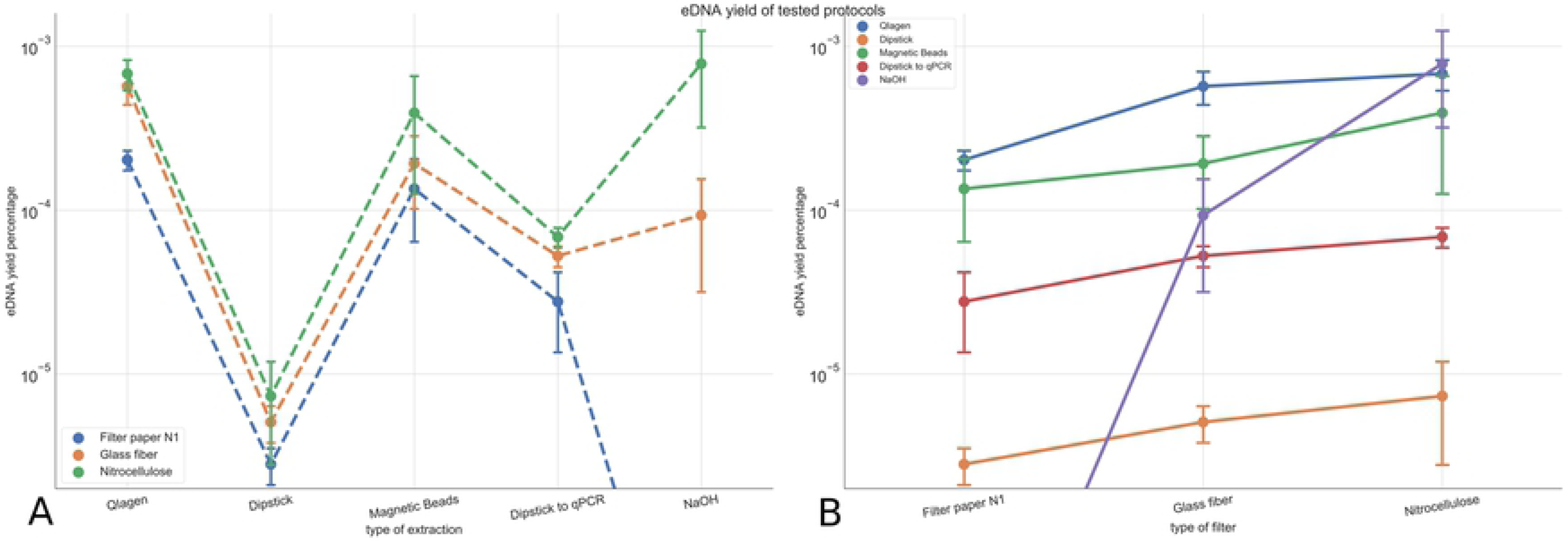
Decision tree for choosing the protocol which will yield the most information given research constraints. A glass filter is recommended in most cases, as long as the focus isn’t maximizing DNA yield with no time or cost constraints. Magnetic beads also are advised in general for its balance between DNA yield and time to process the samples, while cost can be mitigated by producing magnetic beads solution in-house (∼$0.55/mL) instead of buying Ampure XP ($15–$70/mL) [21].

**Figure 8:**
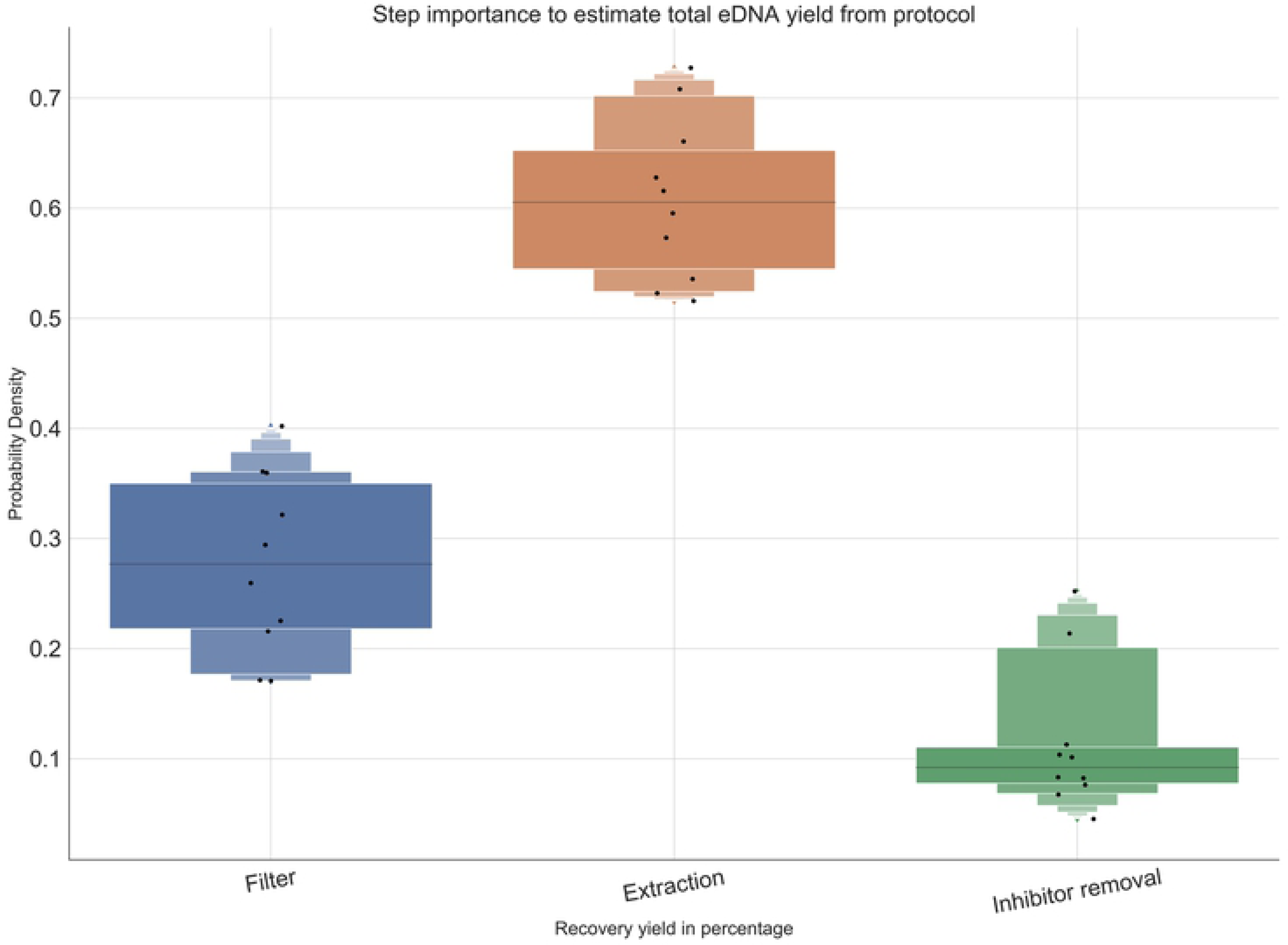
Comparison between eDNA protocols for DNA yield, cost and time to process 96 samples. Methods were sorted by yield and shown in log_10_ scale.

## Methods

### Ethics statement

We sampled water in accordance with the University of California Davis Institutional Animal Care and Use Committee (USDA registration: 93-R-0433, PHS Animal Assurance A3433-01) under the protocol number #20608.

### Experimental Design

We tested three biological replicates for every combination of filter, extraction method and inhibitor removal and measured the amount of recovered eDNA using qPCR Cq values and a DNA standard curve, obtained from a fin clip serial dilution on the same plate (Fig 1). Input DNA estimation is described in the methods section. We defined an equation that describes how the efficiency of each step influences the total amount of recovered eDNA:

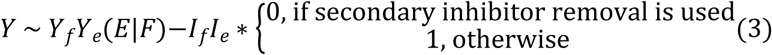

where:

*Y:*ratio of input eDNA that was amplified by the qPCR

*Y*_*f*_:ratio of input eDNA that binds to filter

*Y*_*e*_(*E*|*F*):ratio of eDNA bound to the filter that is isolated by the extraction method

*I*_*f*_:filter inhibitor carryover

*I*_*e*_:extraction method inhibitor carryover

*I*_*f*_*I*_*e*_:ratio of input eDNA not available to amplification due to inhibitors

**Figure 9:**
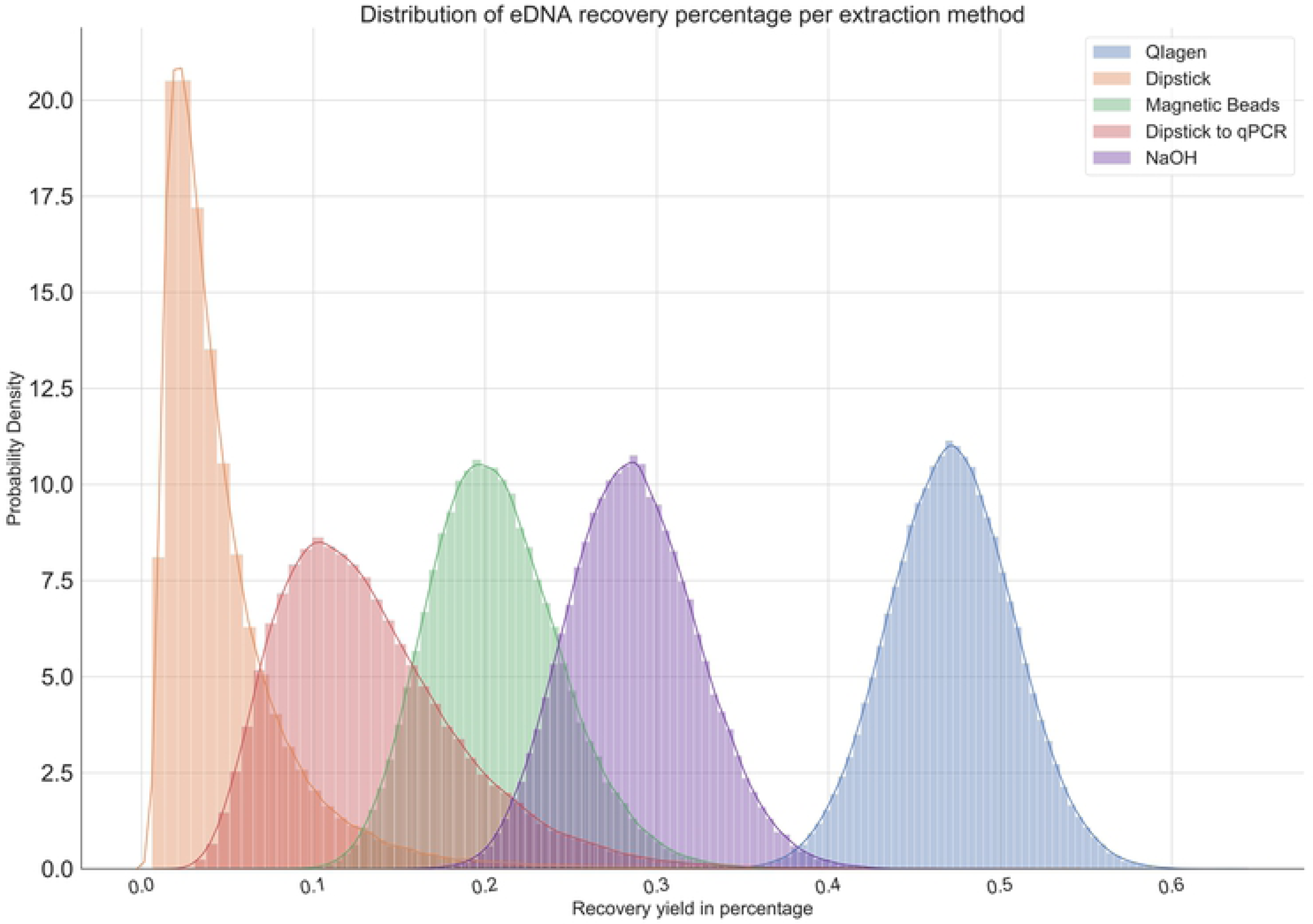
Scheme of steps for an eDNA protocol with tested methods for each step. Cost and processing times (in minutes) of each method are shown next to the method name. Number of samples for the measured times varies as the number of samples that can be run in parallel varies between steps. Costs are estimated per sample.

Then, based on this equation we used Automatic Differentiation Variational Inference (ADVI) [15] to estimate the distribution of the parameters that maximize the likelihood of the observed yields.

### Sampling

To replicate realistic water conditions in terms of salinity, temperature and turbidity while also controlling the presence and amount of Chinook DNA, we combined water samples from a tank containing a high density of juvenile Chinook with an estuarine water sample from a representative location of pre-smolt Chinook habitat in the San Francisco Estuary. The estuarine water biological replicates consisted of 500mL of surface water taken with a 1L measuring cup (sterilized by rinsing in 20% bleach solution and then rinsing in DI water) from Suisun Bay, California (38°11’16.7”N 121°58’34.5”W) and collected in a 1L Nalgene bottle. Next, using another sterile measuring cup, we added 500ml of tank water known to hold Chinook salmon DNA to each estuarine water biological replicate. The 680L tank contained 906 Chinook salmon of approximately 11 cm in length. This mixture allowed us to both control Chinook density and observe similar PCR inhibitor levels as those observed in the estuary. In total, we produced 85 samples, which including a deionized water sample control, a tank water only control and a Suisun Bay water control.

### Estimation of average input eDNA

To estimate the average input DNA from the tank water we spiked 10 samples of 1L surface water from the Suisun Bay, California (38°11’16.7”N 121°58’34.5”W) with varying concentrations of isolated Chinook and green sturgeon (*Acipenser medirostris*) DNA totaling 3 samples with 1ng/L, 3 samples with 0.1ng/L and 3 samples with 0.01ng/L for the Chinook salmon samples and 3 samples with 10ng/L, 3 samples with 1ng/L and 3 samples with 0.1ng/L for green sturgeon. We tested for green sturgeon concomitantly to validate the protocol in a multispecies manner and verify that probe specificity and detection limit doesn’t affect the DNA yield of the protocol. The last sample was not spiked and used as a negative control. Then we filtered the samples using a glass filter, extracted the DNA using the Qiagen DNeasy Blood & Tissue Kit (Cat No./ID: 69504) and removed PCR inhibitors using Zymo OneStep™ (Cat No./ID: D6030). Our serial dilution consisted of the same extracted DNA solution used to spike the samples. We estimated the average yield in percentage for this protocol by qPCR amplification. From the Qiagen protocol average yield we could estimate the average input DNA from the tank water. We also estimated that pipelines DNA yield percentage is mildly inverse correlated (p-value = 0.0023) to the initial DNA concentration (Fig S1), while the probability of amplification is logistically correlated to the Log_10_(initial DNA concentration) (Fig S2).

### Filtration

We filtered the samples one day after sampling to simulate real conditions, where it is not always possible to complete filtration on the same day as sampling. In each filtration run, 4 samples of 1L were filtered in parallel at the speed of 310rpm on a peristaltic pump and we timed each filtration event. Filters were folded in half 3 times and stored in a 2mL microcentrifuge tube and stored at −20°C. Between runs, tubing and casing were sterilized using a bath of 20% bleach [16], rinsed twice using DI water to remove any remaining bleach and dried.

### Extractions

For all DNA extraction protocols except the dipstick-based ones, we added 180µL of ATL buffer and 20µL of 5U Proteinase K to the microcentrifuge tube and incubated at 56°C overnight using a rotisserie attachment for 2mL microcentrifuge tubes. As the incubation time step doesn’t require labor, we didn’t add it to the total time of the protocol. Next, the filter was compressed inside of the microcentrifuge tube using a pipette tip and the supernatant was transferred to a clean 0.5mL (NaOH extraction) or 1.5 mL microcentrifuge tube (magnetic beads and QIagen).

#### NaOH-based extraction

For each 100uL of supernatant we added 5.26µL of 1M NaOH. In a benchtop thermocycler, we incubated the samples at 95°C for 20min and ramped down the temperature at a pace of 0.7°C/min until reaching 4°C. Next, we added 10% of the total volume of 1M Tris-HCL. Samples were vortexed and centrifuged for 15min at 4680rpm. Without disturbing the pellet, 100µL of the supernatant was extracted and transferred to a new 1.5mL microcentrifuge tube and stored at −20°C.

#### Magnetic Beads

For each sample, 180μL of Agencout AMPure XP (Beckman Coulter™; Cat No./ID: A63881) was added to the solution and incubated at room temperature for five minutes. Then the microcentrifuge tubes were placed onto the magnetic plate (DynaMag™-2; Cat No./ID: 123.21D) for 2 minutes. We removed the supernatant and washed the magnetic beads twice using 200μL of a freshly made 70% ethanol solution with an incubation time of 30s in the magnetic plate. We then air dried the beads for 3minutes. A total of 100μL of TE solution was used to resuspend the particles and elute the DNA. The solution was incubated for 1minute at room temperature before pulling down the magnetic beads with the plate for 2minutes. Lastly, the supernatant was transferred to clean 1.5mL microcentrifuge tubes.

#### QIagen DNeasy cell and tissue

The QIagen DNeasy extraction was performed following the manufacturer’s recommendations. A total of 200μL of AL buffer was added and the samples were incubated at 56°C for 10minutes. We added 200μL of ethanol to each sample and the solution was transferred to the column and centrifuged at 8000rpm for 3minutes. Then the column was washed using 500μL of Wash Solution N°1 and centrifuged at 6000rpm for 1minute. Then the column was washed again with Wash Solution N°2 and centrifuged for 3minutes at 1400rpm. Next 100μL of AE solution was added and incubated for 20minutes before centrifuging at 8000rpm for one minute. The flowthrough was then stored at −20°C

#### Whatman paper dipstick

Dipsticks were made following the protocol described in [17]. We used the qualitative Whatman filter n°1 to make our dipsticks and used an effective surface area of 8mm^2^ (2mm width and 4mm height). We added 200μL of lysis buffer and ground the filter using a pipette tip until the filter was dissolved. Then we dipped the dipstick in the lysis buffer solution (20mM Tris [pH 8.0], 25mM NaCl, 2.5mM EDTA, 0.05% SDS) 3 times, then dipped 3 times in 100μL of wash solution (10mM Tris [pH 8.0], 0.1% Tween-20), and 3 times in a final solution of nuclease free water which then was stored at −20°C. In the case of “straight to qPCR” dipstick extraction, we directly dipped the dipstick after the wash step into the qPCR reaction.

### Secondary Inhibitor removal

Zymo OneStep™ PCR Inhibitor Removal Kit (Zymo Research; Cat No./ID: D6030) was used following the manufacturer’s protocol in order to remove any carryover PCR inhibitors from previous steps. We added 600μL of Prep-solution to the column and centrifuged at 8000g for 3 minutes, the flow through was discarded, then 50μL of DNA elute from previous steps were added to the column and centrifuged at 16000g. The flowthrough was then stored at −20°C.

### qPCR amplification oligos

Quantitative PCR detection for Chinook was developed by adapting the protocol from [18]Reaction solution totaling 20μL was composed of 1× TaqMan™ Environmental Master Mix 2.0 (ThermoFisher Scientific; Cat No./ID: 4396838), 0.9µM concentration of each primer, and 0.7µM of the Taqman probe, and 6µl isolated DNA extract from previous steps. Thermocycling was performed on a Bio-Rad CFX96 real-time detector using the following profile: 10 min at 95°C, 40 cycles of 15s denaturation at 95°C and 1 min annealing–extension at 60°C.

**Table 1:**
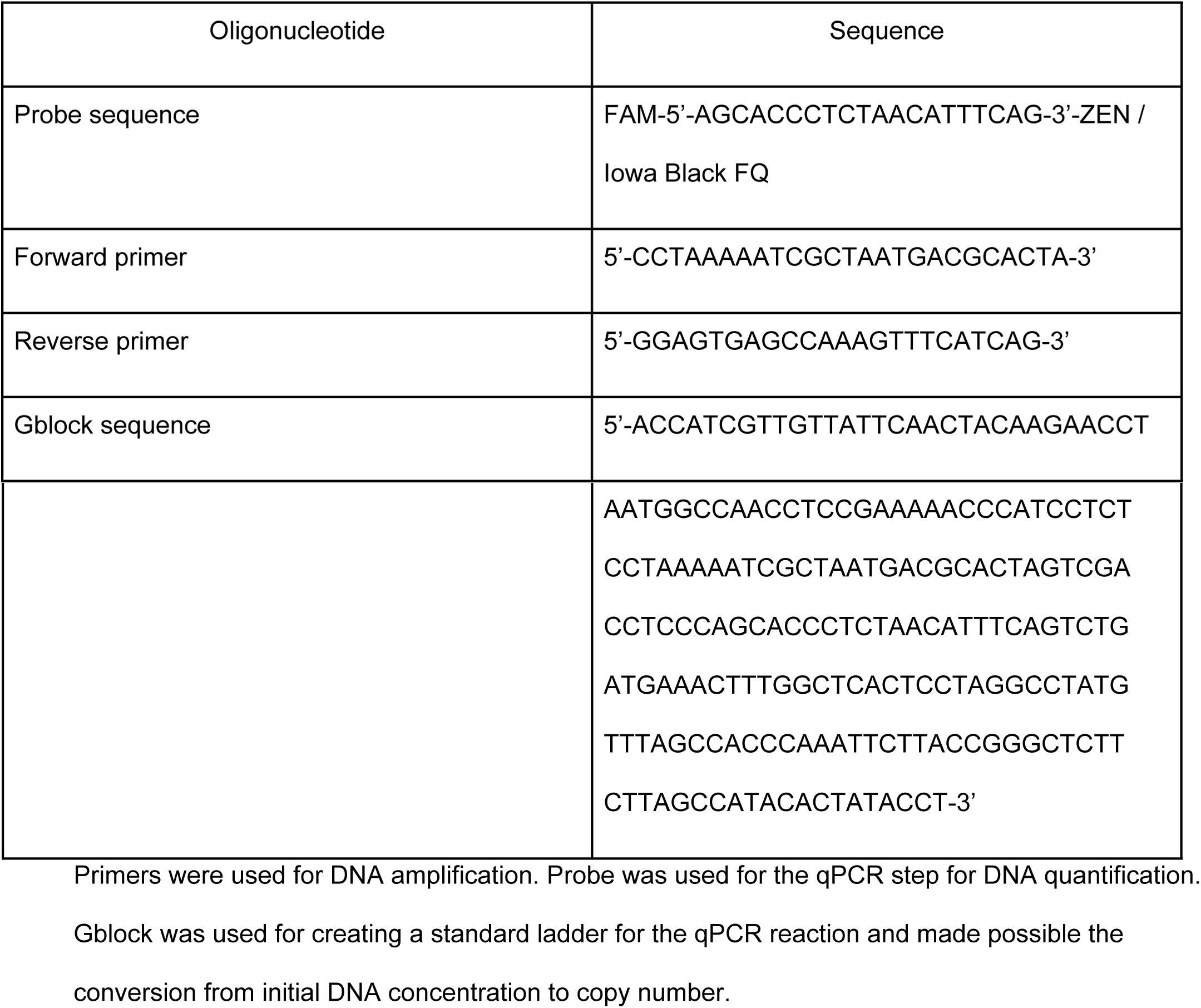
List of used DNA oligonucleotides.

## Data analysis

Data analysis was performed in Python 3.7 and is available on https://github.com/sanchestm/eDNA-Protocol-Optimization. We measured interference between filter type and extraction method using two competing models, one that includes the interference effect and one that does not. Using ADVI inference we fitted the data to the models [15]. From the ADVI fitting for the best model we estimated the distribution of filter eDNA yield percentage (Fig 2), extraction eDNA yield percentage (Fig 4) and PCR inhibitor carryover for filtration and extraction (Fig 6). To estimate which step of an eDNA experiment has the most variance between methods, and therefore can lead to the most significant gains when optimized, we trained a random forest regressor [19] with the collected data and estimated importance of each step of the experiment.

<u>PHS/NIH Assurance (A3433-01)</u>

## Supporting information

**Figure S1:**
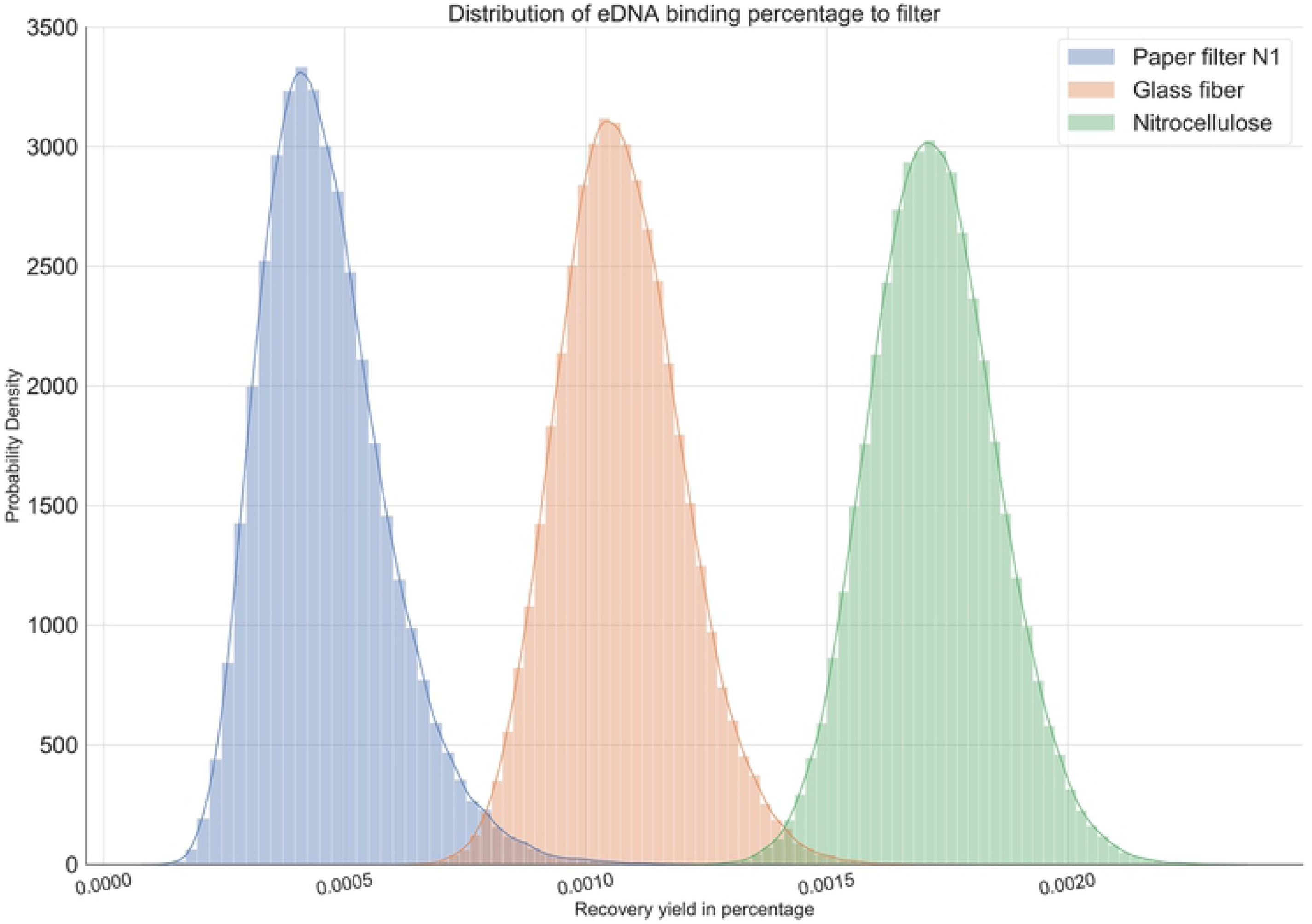
Correlation between DNA yield and initial eDNA concentration. Blue dots - median value; vertical lines - 95% CI; horizontal line - linear regression between protocol DNA yield and input DNA, with both axes being represented in log_10_ scale. P(amp) - probability of amplification.

**Figure S2:**
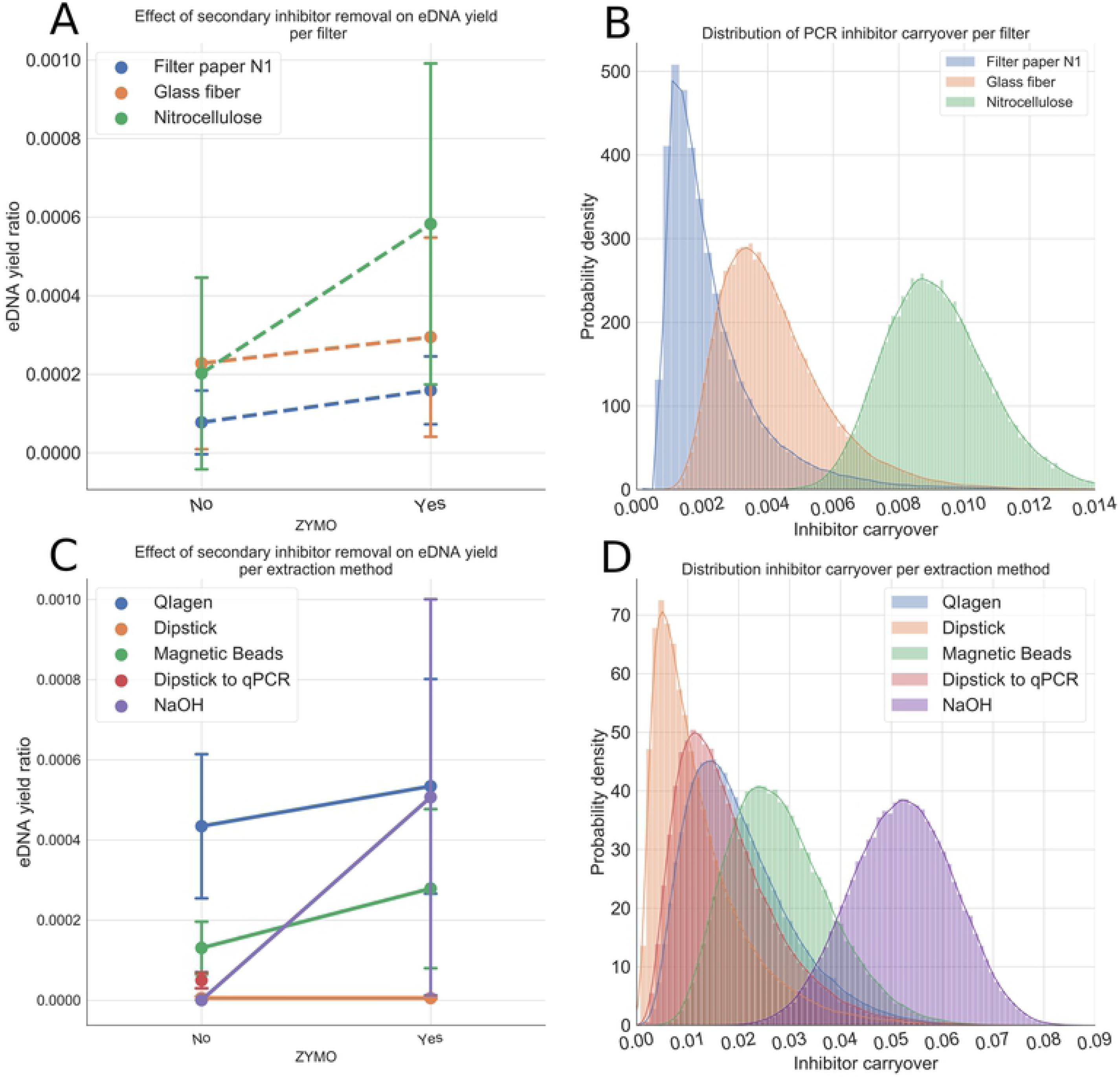
Probability of amplification as a function of the input DNA concentration. Dots - probability of amplification from DNA spiking experiment; line - logistic fit to data points.

